# Antigen B from *Echinococcus granulosus* enters mammalian cells by endocytic pathways

**DOI:** 10.1101/224691

**Authors:** Edileuza Danieli da Silva, Martin Cancela, Karina Mariante Monteiro, Henrique Bunselmeyer Ferreira, Arnaldo Zaha

## Abstract

Cystic hydatid disease is a zoonosis caused by the larval stage (hydatid cyst) of *Echinococcus granulosus* (Cestoda, Taeniidae). The hydatid cyst develops in the viscera of intermediate host as a unilocular structure filled by the hydatid fluid, which contains parasitic excretory/secretory products. Antigen B (AgB) is the major component of *E. granulosus* metacestode hydatid fluid. Functionally, AgB has been implicated in immunomodulation and lipid transport. However, the mechanisms underlying AgB functions are not completely known. In this study, we investigated AgB interactions with different mammalian cell types and the pathways involved in its internalization. AgB uptake was observed in four different cell lines, NIH-3T3, A549, J774 and RH. Inhibition of raft-mediated endocytosis causes about 50 and 69% decrease in AgB internalization by RH and A549 cells, respectively. Interestingly, AgB colocalized with the raft endocytic marker, but also showed a partial colocalization with the clathrin endocytic marker. The results indicate that raft-mediated endocytosis is the main route to AgB internalization, and that a clathrin-mediated entry may also occur at a lower frequency. Cellular internalization could be a requirement for AgB functions as a lipid carrier and/or immunomodulatory molecule, contributing to create a more permissive microenvironment to metacestode development and survival.

**Author summary:** Antigen B (AgB) is an oligomeric lipoprotein highly abundant in *Echinococcus granulosus* hydatid fluid. AgB has already been characterized as an immunomodulatory protein, capable of inducing a permissive immune response to parasite development. Also, an important role in lipid acquisition is attributed to AgB, because it has been found associated to different classes of host lipids. However, the mechanisms of interaction employed by AgB to perform its functions remain undetermined. In this study, we demonstrate that mammalian cells are able to internalize *E. granulosus* AgB in culture and found that specific mechanisms of endocytosis are involved. Our results extend the understanding of AgB biological role indicating cellular internalization as a mechanism of interaction, which in turn, may represent a target to intervention.

## Introduction

Cystic hydatid disease (CHD), caused by the larval stage (hydatid cyst or metacestode) of parasites belonging to the *Echinococcus granulosus sensu lato* (s.l.) complex, is a zoonosis of worldwide occurrence, with a considerable medical and economic impact [1]. CHD is endemic or hyperendemic in South America, especially in Argentina, Southern Brazil, Uruguay, Chile and mountainous regions of Peru and Bolivia [2]. In 2010, the World Health Organization added CHD to its list of Neglected Tropical Diseases (http://www.who.int/neglected_diseases/diseases/en/). *Echinococcus granulosus sensu stricto*, or simply *Echinococcus granulosus*, is one of the cryptic species of the *E. granulosus* s.l. complex and is the species most widely distributed worldwide. Also, *E. granulosus* is responsible for most cases of human CHD infections [3].

The adult tapeworm lives in the small intestine of a definitive canid host, and the larval stage develops in the viscera of a wide range of mammal species, including humans. *E. granulosus* life cycle is predominantly domestic, where dogs are the definitive hosts and ungulates are the intermediate hosts [4]. The metacestode is a fluid-filled, unilocular cyst containing protoescoleces in its lumen. Protoescoleces are the pre-adults, infective to the definitive host, which remain quiescent and immersed in the hydatid fluid (HF), which is a complex mixture of molecules of both host and parasite origin. The excretory/secretory products of the metacestode are of special relevance for the host-parasite relationship, as they have a greater potential to interact with host proteins and cells.

Antigen B (AgB) is the most abundant and the major immunodominant protein among the excretory/secretory metacestode products in the HF. AgB belongs to the group of hydrophobic ligand binding proteins (HLBPs), a cestode protein family whose members are known by their high abundance and immunogenicity, and by their oligomeric structure, comprising 7-10 kDa α-helix rich subunits [5,6]. The AgB oligomeric structure comprises 8 kDa subunits (AgB8/1 to AgB8/5) encoded by a multigene family [7], which are differentially expressed among the parasite life-cycle stages, metacestode tissues and individuals [8–10]. AgB oligomers have been observed predominantly in the molecular mass range of 150-200 kDa, but aggregates with higher molecular masses have also been detected [9,11].

It has been demonstrated that delipidated AgB is able to bind hydrophobic compounds *in vitro* [12]. The lipid moiety associated with AgB was analyzed and different lipids were identified, with cholesterol, phospholipids and triacylglycerides being the most prominent [13]. Moreover, delipidated recombinant AgB8/2 and AgB8/3 subunits were capable of transferring fatty acids analogues to artificial phospholipid membranes [14]. *E. granulosus* genome lacks sequences for several key enzymes for fatty acid and cholesterol synthesis, thus the parasite is incapable of synthesizing these compounds *de novo* [15,16]. Hydatid cyst viability relies on the sequestration and utilization of host lipids, and AgB might be involved in lipid uptake from host tissue and its transport to the parasite, by stabilizing insoluble lipids into a lipoproteic particle [13].

In addition, AgB roles in the modulation of both innate and adaptive immunity have been proposed. It has been described that neutrophils have both the recruitment inhibited and hydrogen peroxide production decreased by AgB [17,18]. Besides, AgB polarizes the immunological response to a Th2 profile, which is protective to the parasite [19,20].

Considering the two main roles attributed to AgB, immunomodulation and lipid transport, it is reasonable to consider that a direct interaction with host cells and tissues should occur. In fact, it was recently demonstrated that AgB binds to macrophages and the plasma membrane of inflammatory monocytes, inducing a non-inflammatory phenotype in macrophages [21]. However, little is known about the molecular details of AgB-cell interaction and whether AgB interacts with non-immune cells, or even enters into the cell.

In the present work, we investigated the ability of HF-purified AgB oligomers to enter into different mammalian cell types *in vitro*, and the mechanisms involved in AgB internalization. Immunopurified AgB was incubated with four distinct cell lines representative of different cell types, namely hepatocytes, fibroblasts, macrophages, and lung epithelial cells. We demonstrated the entry of AgB into the cytoplasm of all studied cell lines. Moreover, we provided evidence that the endocytic pathways are involved in AgB internalization by cells, with raft-mediated endocytosis being the prevailing one.

## Methods

### Biological material

Bovine viscera containing hydatid cysts from *E. granulosus* were obtained from a local slaughterhouse (São Leopoldo, Brazil). Animal slaughtering was conducted according to Brazilian laws and under supervision of the *Serviço de Inspeção Federal* (Brazilian Sanitary Authority) of the Brazilian *Ministério da Agricultura, Pecuária e Abastecimento*. HF was removed by punction and aspiration from individual fertile cysts and kept at −80°C until use. Parasite genotyping was performed for species determination [22].

### Immunoblot

Aliquots of 100 μl of HF samples were resolved on SDS-PAGE 12% and electrophoretically transferred onto a nitrocellulose membrane. A pool of rabbit polyclonal antibodies raised against each recombinant AgB subunit (AgB8/1 to 5) were used at 1:70.000 dilution as primary antibody. A horseradish peroxidase-conjugated goat anti rabbit IgG (GE Healthcare) diluted at 1:7.000 was used as the secondary antibody. Blots were developed using the chemiluminescent reagent ECL Plus (Pierce, ThermoScientific) and imaged in VersaDoc system (BioRad). HF samples with higher AgB content were used for the protein purification step (S1A Fig).

### *E. granulosus* AgB purification

AgB purification was carried out following the protocol described by Oriol *et al*. [23], with some modifications. Briefly, parasite proteins from *E. granulosus* HF were precipitated by sodium acetate (5 mM, pH 5.0) and the resultant material was resuspended in phosphate-buffered saline (PBS) containing 20 μM 3,5-di-tert-butyl–4-hydroxytoluene (BHT). The HF parasite enriched fraction was subjected to immunoaffinity chromatography using rabbit polyclonal antibodies against the recombinant forms of AgB8/1, AgB8/2 and AgB8/4. Antibodies were separately coupled to cyanogen bromide-activated Sepharose™ 4B resin (GE Healthcare) and the previously prepared HF material was passed through the columns. Bound AgB from each column was eluted with 100 mM tris-glycine pH 2.5, then pooled together, dialyzed against PBS/BHT and concentrated on Amicon Ultra-15 centrifugal filter device, MWCO 3 kDa (Millipore). Purified AgB was analyzed on SDS-PAGE 12% (S1B Fig). AgB concentration was determined using a Qubit quantitation fluorometer and Quant-iT reagents (Life Technologies).

### Cell cultures

NIH-3T3 (mouse fibroblasts), A549 (human lung adenocarcinoma), J774 (mouse macrophages) and RH (rat hepatoma) cells were cultivated in DMEM containing 10% fetal bovine serum, 100 U/ml penicillin and 100 μg/ml streptomycin in a 5% CO_2_ humidified environment at 37°C. J774 culture media was also supplemented with MEM non-essential amino acid solution, 2 mM glutamine, 10 mM HEPES and 1 mM sodium pyruvate. All cells lines were free from mycoplasma contamination.

### AgB internalization assays

NIH-3T3, A549, J774 and RH cells were grown on sterile glass coverslips in 35 mm Petri dishes. Cell media was changed to serum-free medium and the cells were then incubated with 40 μg/ml of AgB oligomers for 4 h at 37°C, or 4°C. Controls were incubated with equal volume of PBS/BHT. Unbound protein was then removed by three washes with cold PBS and cells were fixed in 4% paraformaldehyde/PBS at room temperature for 15 min.

In all microscopy preparations, a pool with the same proportion of polyclonal antibodies against AgB8/1, AgB8/2 and AgB8/4 subunits was used as primary antibody for detection of AgB oligomers. Fixed cells were permeabilized with 0.2% Triton X-100/PBS and unspecific sites were blocked with 5% BSA in PBS-T (PBS with 0.05% Tween-20). After, cells were incubated overnight at 4°C with the primary antibodies (1:500) and then for 1 h with 1:200 diluted Alexa Fluor^®^ 488-conjugated anti-rabbit secondary antibody (Molecular Probes) at room temperature. Nuclei were stained with 100 nM 4’,6-diamidino-2-phenylindole (DAPI) (Molecular Probes). Actin was stained with 50 nM Alexa Fluor^®^ 594-conjugated phalloidin (Molecular Probes). Cells were imaged using a LSM 710 Zeiss confocal microscope.

The fluorophore CM-DiI (Molecular Probes) was used to directly label AgB oligomers, because it has affinity to the lipidic compounds associated to the protein. DiI-labelled AgB was used to analyze internalization without cells fixation. AgB was labelled with 5 μM CM-DiI (Molecular Probes) for 1 h at room temperature. Dye excess was washing out with 5-fold the original PBS volume on Amicon Ultra-0.5 centrifugal filter devices, NMWL 100 kDa (Millipore). RH cells were incubated with 40 μg/ml of DiI-labelled AgB oligomers for 4 h at 37°C, washed three times with cold PBS, and immediately analyzed using an Olympus FluoView 1000 confocal microscope.

### Endocytosis inhibition assays

RH and A549 cell monolayers were grown on sterile glass coverslips in six-well tissue culture plates. After changing the cell media to serum-free DMEM, cells were pretreated with endocytosis inhibitors for 30 min at 37°C. A pilot test, where cells were incubated with different concentrations of the inhibitors, was conducted to determine the best concentration to be used. The highest concentration where >80% of the cells remained attached and with little morphological alterations was chosen.

Genistein (Santa Cruz Biotechnology) was used at 100 μg/ml concentration and chlorpormazine (Santa Cruz Biotechnology) at 5 μg/ml. AgB was then added at 40 μg/ml and after incubation at 37°C for 1.5 h, the unbound proteins were removed by acidic stripping (0.5 M NaCl, 0.5% acetic acid, pH 3.0) and three washes with cold PBS. Cells were fixed and prepared for microscopy as described above. Cells were imaged using an Olympus FluoView 1000 confocal microscope. Immunofluorescence intensity normalized by cell area was assessed with ImageJ software [24]. Image analysis was done on two (A549) or three (RH) independent experiments, where three microscopy fields were counted for each experiment (100–300 cells/experiment).

### Colocalization assays

RH cell monolayers were grown on sterile glass coverslips in 6-well tissue culture plates. Cell media was replaced to serum-free DMEM containing 40 μg/ml AgB and the distribution of internalized protein was compared with that of different endocytic markers following up to 1.5 h incubation at 37°C. Endogenous transferrin receptors were labeled with 50 μg/ml Alexa Fluor^®^ 633-conjugated transferrin (Tfn) (Molecular Probes), added in the last 45 min. Alexa Fluor^®^ 555-conjugated cholera toxin subunit B (Ctx-B) (Molecular Probes) at 1 μg/ml concentration was added in the last 15 min of incubation. Adsorbed and unbound proteins were removed by acidic stripping (0.5 M NaCl, 0.5% acetic acid, pH 3.0) and three washes with cold PBS. Cells were prepared for microscopy and imaged as described for endocytosis assay. Colocalization was assessed using JaCoP plugin from ImageJ software [25]. Image analysis was done for two independent experiments.

### MTT reduction assays

A549 and RH cells were plated onto 96-well plates at a density of 10^4^ cells/well. AgB oligomers were added to the cell media at 2.5 – 40 μg/ml final concentrations. After 24 h incubation, 0.5 mg/ml of MTT solution in PBS was added to each well and incubated for a further 4 h. To solubilize formazan, 100 μl of cell lysis buffer (16% SDS, 40% N,N-dimethylformamide, 2% acetic acid, pH 4.7) was added to each well and the samples were incubated overnight at 37°C in a humidified incubator. Absorbance values of formazan were determined at 595 nm with an automatic microplate reader (Bio-Rad, model 550). Analysis was done for five independent experiments.

### Statistics

A Kolmogorov-Smirnov was applied to verify the normality of the data. Statistical significance was analyzed by unpaired Student’s t-test using the GraphPad Prism 6.0 software. Data are expressed as mean ± SEM and p values of less than 0.05 were considered statistically significant.

## Results

### AgB oligomers are internalized by mammalian cells in culture

To investigate the ability of AgB oligomers to interact with and to be internalized by mammalian cells, AgB from *E. granulosus* HF was immunopurified and added to the culture medium of NIH-3T3, A549, RH or J774 cells. AgB internalization was evaluated after 4 h of incubation at 37°C using an immunofluorescence assay. Cells were prepared for confocal microscopy by labelling AgB oligomers with polyclonal antibodies against AgB8/1, 2 and 4 subunits and a secondary anti-rabbit IgG conjugated to Alexa Fluor^®^ 488. AgB signals were detected in the four cell lines tested, suggesting that AgB is able to interact with mammalian cells by a mechanism independent of cellular type. No signals were detected in the cells without AgB (Fig 1).

**Fig. 1.**
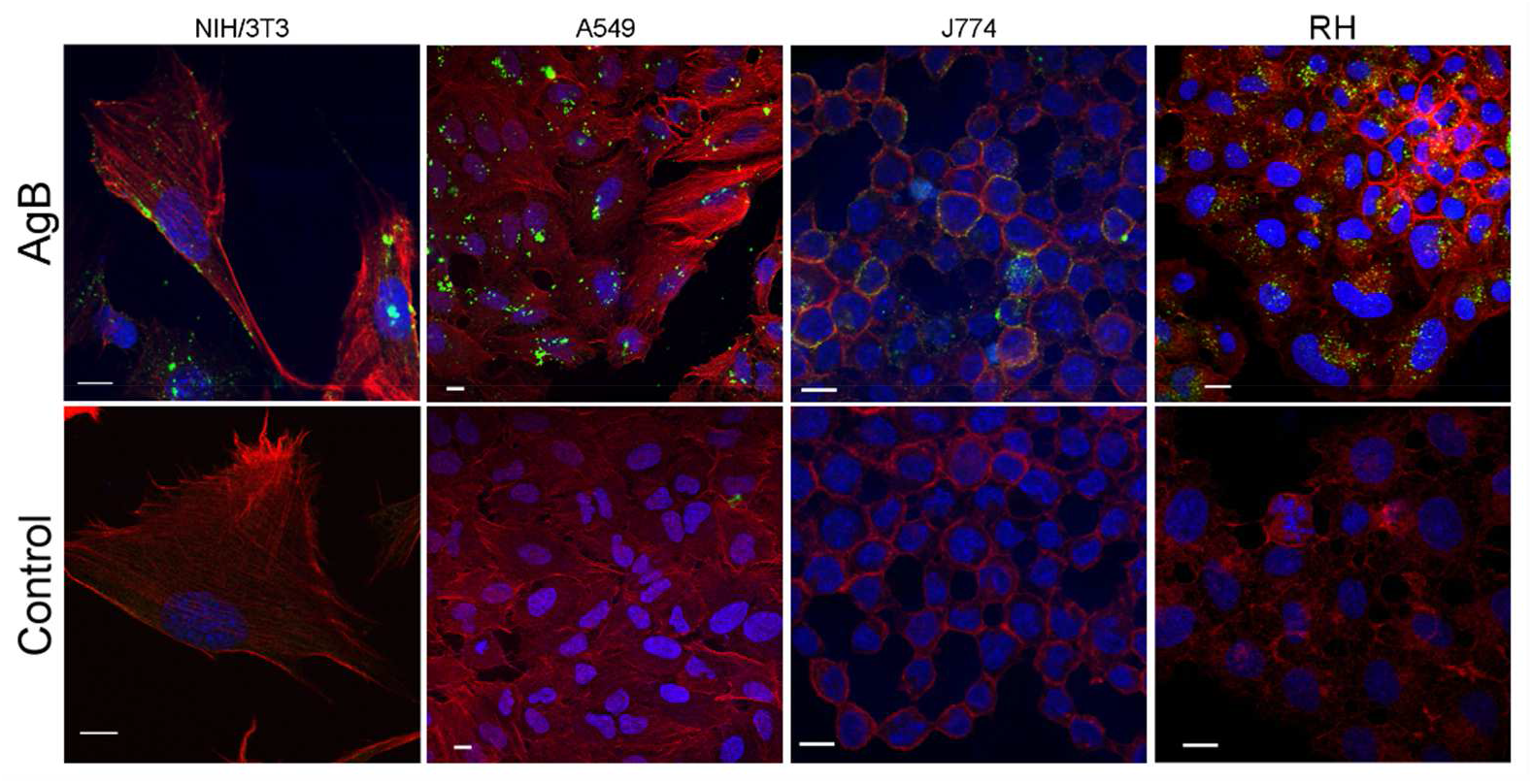
*E. granulosus* AgB uptake by mammalian cells in culture. Immunofluorescence assay was performed on NIH-3T3, A549, J774 and RH cells exposed to 40 μg/ml AgB for 4 h, and mock treated cells (Control). AgB was labeled with antibodies against AgB8/1, 2 and 4 subunits and an Alexa Fluor^®^ 488-conjugated secondary antibody (green). Nuclei and cytoskeleton were stained with DAPI (blue) and Alexa Fluor^®^ 594-conjugated phalloidin (red), respectively. Images are median optical sections from z-stacks obtained by confocal microscopy. Scale bar, 10 μm.

To confirm the internal localization of AgB in the cells, the oligomers were labelled with DiI and incubated with RH cells in the same way as before. However, the analysis on confocal microscope was conducted right after incubation had been finished, without cell fixation. The intermediate sections from confocal z-stacks showed higher AgB signal than top or bottom sections, confirming that AgB was inside cells and not just adsorbed to cell membrane (S2 Fig).

AgB was detected in the cell cytoplasm, but not in the nucleus. In addition, Fig 1 and S2 Fig show vesicular-like distribution of AgB oligomers in the cytoplasm of the cell lines analyzed, indicating an internalization through endocytosis. Supporting this idea, AgB internalization does not occur when RH cells were incubated at low temperature (S3 Fig), a condition known to interfere in endocytosis-dependent cellular internalization [26].

### Endocytic pathways involved in AgB internalization

Having established that AgB oligomers could access the cytoplasmic compartment of mammalian cells, we then investigated which endocytic pathway could be responsible for this AgB uptake by RH and A549 cells. These two cell lines were chosen to perform the following experiments in an attempt to simulate the natural situation, as liver and lungs are the primary organs infected by *E. granulosus*.

Genistein, a tyrosine kinase inhibitor that prevents lipid raft-mediated endocytosis, was used to treat RH cells; and after 30 min AgB oligomers were added to the culture media and left to incubate for another 1.5 h at 37°C. We found that internalization of AgB was inhibited by ∼50% in RH cells treated with genistein (Figs 2A and 2B). The same inhibition assay was carried out with A549 cells and we found very similar results, where AgB uptake was inhibited by ∼69% (Figs 2A and 2B). The results were statistically significant for both cell lines (Fig 2B).

**Fig. 2.**
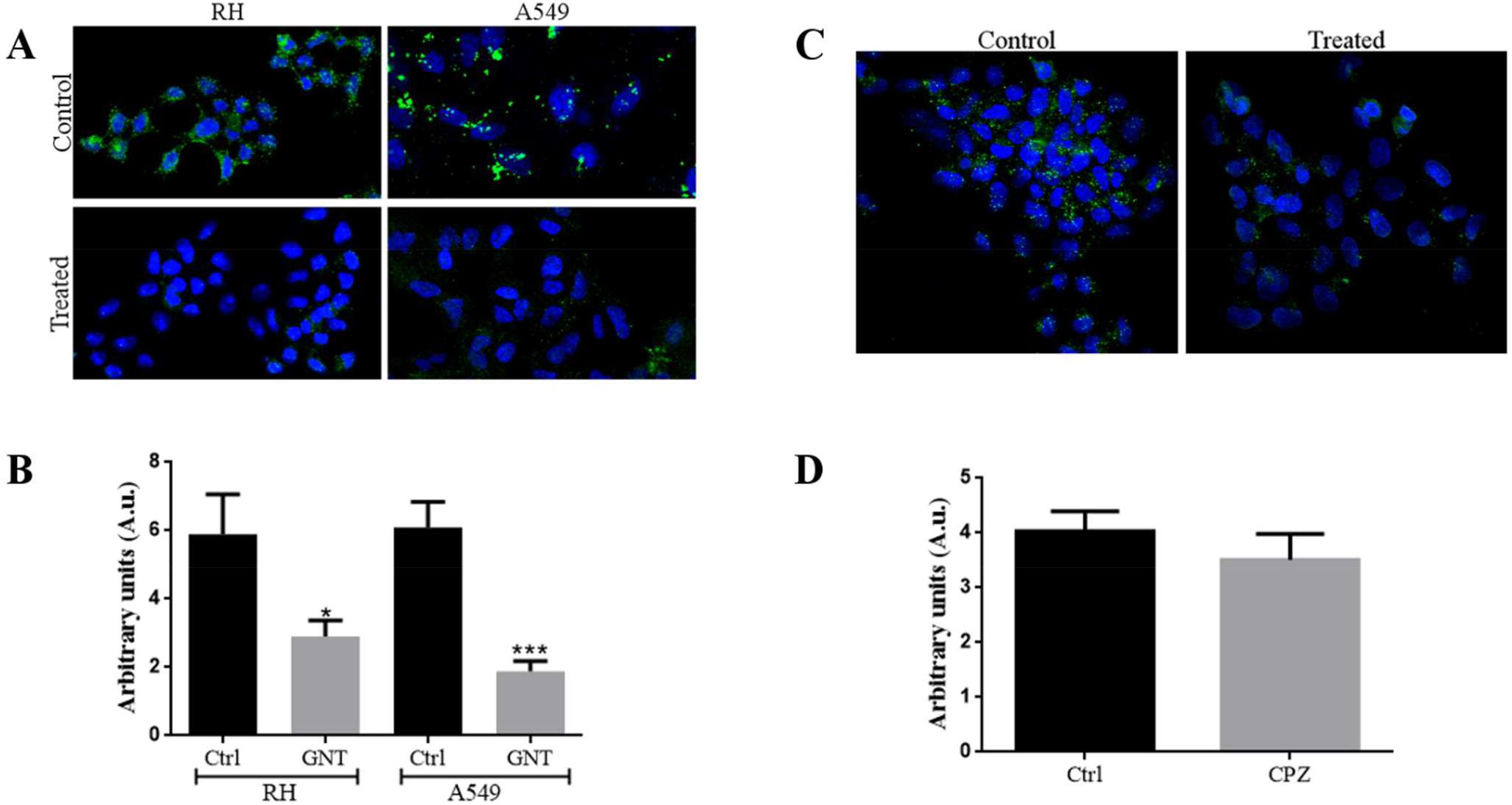
Raft-mediated endocytosis is the main route involved in *E. granulosus* AgB internalization by A549 and RH cells. *A*, inhibition of raft-mediated endocytosis by genistein reduces AgB internalization. *Lower panels*, RH and A549 cells pre-treated with 100 μg/ml genistein, then exposed to 40 μg/ml AgB for 1.5 h. *Upper panels*, non-treated cells. *B*, quantitative data for AgB internalization after genistein (GNT) treatment represented in *A. C*, inhibition of clathrin-mediated endocytosis pathway by chlorpromazine does not cause a significant decrease in uptake of AgB by RH cells. *Right panel*, cells treated with 5 μg/ml chlorpromazine for 30 min, then exposed to 40 μg/ml AgB for 1.5 h. *Left panel*, non-treated cells. *D*, quantitative data for AgB internalization after chlorpromazine (CPZ) treatments represented in *B*. AgB was detected using antibodies against AgB8/1, 2 and 4 subunits and a secondary anti-rabbit IgG Alexa Fluor^®^ 488 conjugated antibody (green). Cell nuclei were labeled with DAPI (blue). Ctrl: control. Measurements from three experiments with RH cells and two with A549 cells were averaged. Error bars indicate SEM. *p=0.037, ***p=0.0004 according to Student’s t-test.

In order to test whether other endocytic pathways could be involved in AgB oligomers uptake by RH cells, the inhibition assay was performed using chlorpromazine, an inhibitor of clathrin-mediated endocytosis. After chlorpromazine treatment, internalization of AgB was reduced by ∼13%, however this was not statistically significant (p = 0.39) (Figs 2C and 2D). The higher inhibition of AgB internalization in the genistein treatment indicates that raft-mediated endocytosis is the major pathway associated with AgB uptake.

The specificity of the inhibitors was confirmed using established endocytic markers as controls. Internalization of transferrin (Tfn), which undergoes clathrin-mediated endocytosis, was affected only by chlorpromazine. In contrast, genistein only reduced internalization of cholera toxin subunit B (Ctx-B), which undergoes raft-mediated endocytosis (S4 Fig).

In a complementary approach, the distribution of internalized AgB was compared with that of established endocytic markers, Tfn and Ctx-B. RH cells were cultivated in the presence of AgB for 1.5 h, with either Tfn or Ctx-B being added in the last 45 and 15 min of incubation, respectively. Results were analyzed by confocal microscopy and the level of colocalization between the two fluorophores and, consequently, the two proteins, was determined according to Pearson’s correlation coefficient. The results indicated that AgB was colocalized with Ctx-B (Pearson’s coefficient = 0.64 ± 0.02) (Fig 3), which is in accordance with our previous results that raft-mediated endocytosis is involved in AgB uptake. Interestingly, AgB seemed partially colocalized with Tfn (Pearson’s coefficient = 0.49 ± 0.04), suggesting that in some degree AgB could be internalized by clathrin-mediated endocytosis.

**Fig. 3.**
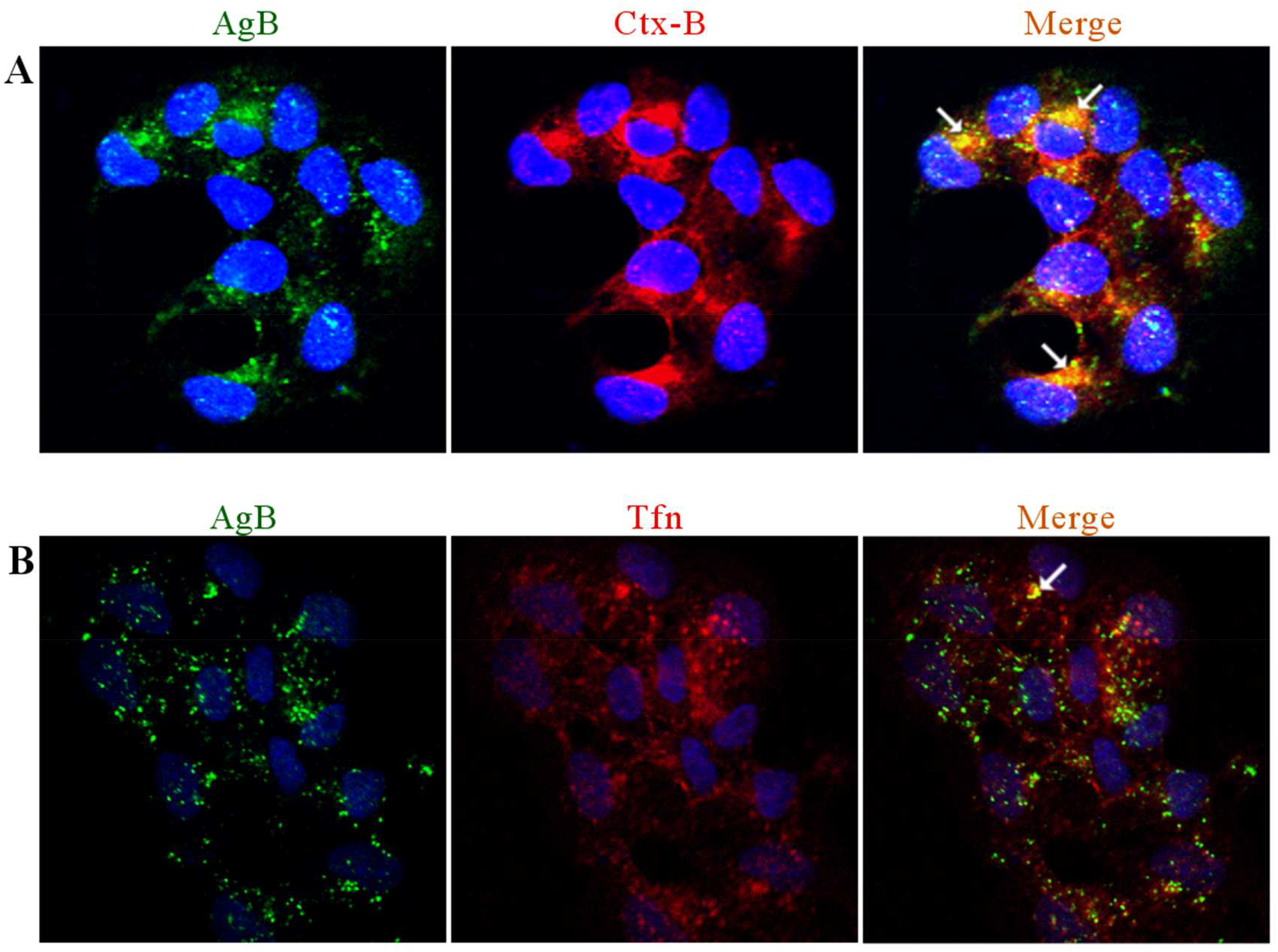
Internalized *E. granulosus* AgB colocalizes with protein endocytic markers in RH cells. Alexa Fluor^®^ 633-conjugated Tfn was used as a marker of clathrin-mediated endocytosis and Alexa Fluor^®^ 555-conjugated Ctx-B was used as a marker of raft-mediated endocytosis. Confocal microscopy images of RH cells incubated with 40 μg/ml AgB and 1 μg/ml Ctx-B (A) or 50 μg/ml Tfn (B) are presented. AgB was detected using antibodies against AgB8/1, 2 and 4 subunits and a secondary anti-rabbit IgG Alexa Fluor^®^ 488 conjugated antibody (green). Arrows indicate colocalization points. Cell nuclei were labeled with DAPI (blue). The endocytic markers are shown in red.

Altogether, the above findings provide evidence that AgB entry into mammalian cells occurs mainly via raft-mediated endocytosis, although it could also occur by clathrin-mediated endocytosis in a lesser extent.

### AgB oligomers do not induce cellular toxicity

To further investigate the possible effects of AgB oligomers internalization by mammalian cells, A549 and RH cells were incubated with different concentrations of AgB (2.5-40 μg/ml) for 24 h. Alterations of cell physiologic state were then evaluated by MTT reduction assays. Both cell lines did not present any significant decrease in their ability to metabolize MTT in presence of AgB oligomers (Fig 4). AgB oligomers do not have a cytotoxic effect on cells, so this interaction is probably part of the mechanisms underlying AgB function in host-parasite interplay.

**Fig. 4.**
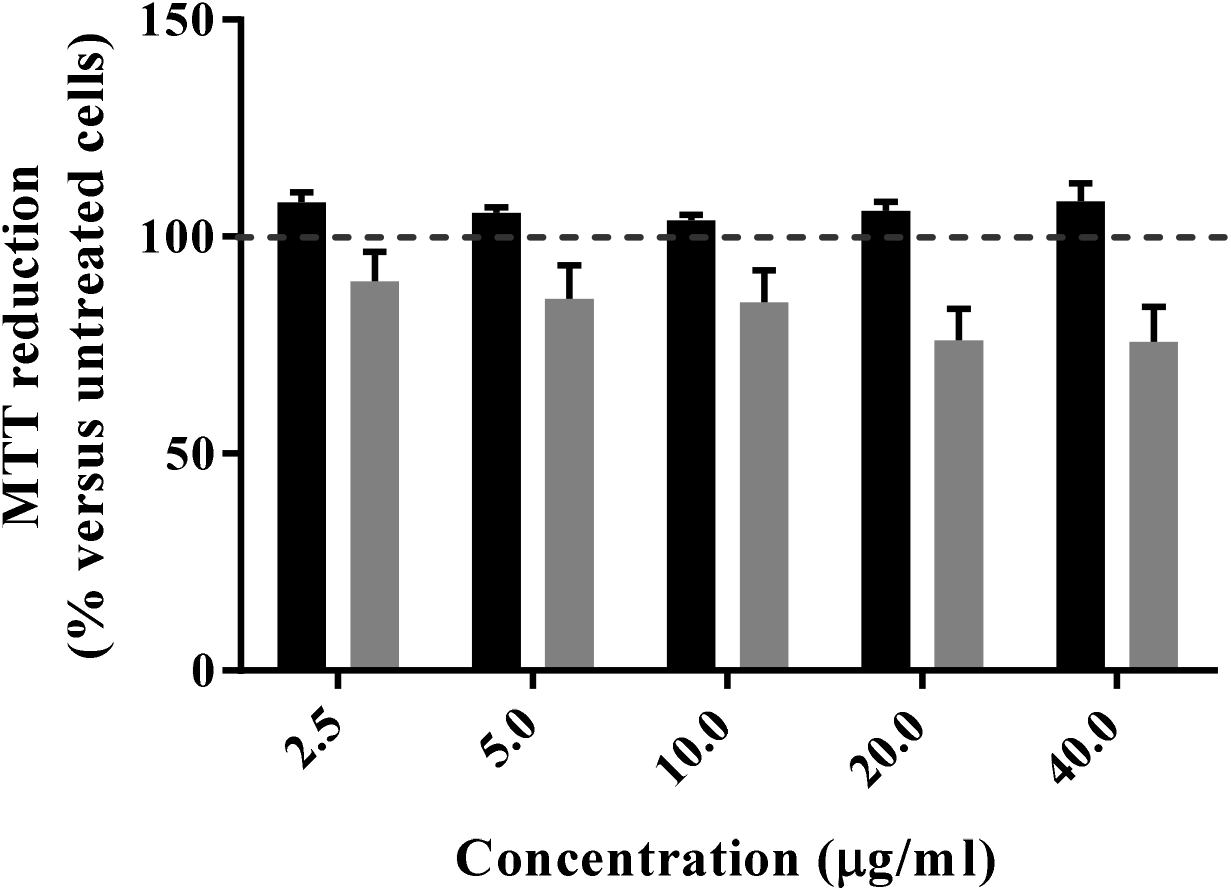
MTT reduction assay. A549 (black bars) and RH (gray bars) cells were treated for 24 h with the indicated concentrations of *E. granulosus* AgB oligomers. Cell viability is expressed as percentage of MTT reduction measured for untreated cells and assumed as 100% (horizontal dashed line). Error bars correspond to the SEM values of five independent experiments.

## Discussion

A long-term growth is characteristic of the chronic infection caused by *E. granulosus* metacestodes, and to allow that, different molecular mechanisms are employed by the parasite to ensure its survival and development in the host microenvironment. In an attempt to evade the host’s immune response and absorb nutrients from the host, the parasite secretes several molecules. AgB is the major antigen in HF and has been implicated in immunomodulation processes, as well as in lipid transport and uptake [13,27].

AgB has been studied for a long time and it is known to generate a strong humoral response and to modulate host immune response, which is in agreement with the idea that AgB is secreted towards the outside of the hydatid cyst [27,28]. Moreover, there are many studies reporting AgB effects on immune cells that support a direct AgB-cell interaction in host tissue [20,21]. However, whether this interaction occurs with other cell types and what underlying mechanisms are involved in this process, are still unclear. We hypothesized that AgB may interact with host tissue cells surrounding the hydatid cyst to interfere with host cell homeostasis, facilitating nutrient acquisition and immune evasion. In this study, we demonstrated that *E. granulosus* AgB is taken up by mammalian cells *in vitro* by endocytosis. Since AgB internalization seems to be independent of the cell type, it most likely occurs by a ubiquitous mechanism. Thus, we further evaluate the involvement of specific endocytic pathways on AgB internalization.

Clathrin-mediated endocytosis is the best-understood internalization pathway and refers to intake of receptors and their bound ligands through vesicles which are coated by the protein clathrin [29]. Among the molecules known to be internalized by this pathway is the Tfn receptor, hence Tfn was used as a marker for clathrin-mediated endocytosis [30]. Besides the clathrin-mediated pathway, lipid rafts domains are important contributors to endocytosis processes. These are heterogeneous membrane domains enriched in sphingolipids and cholesterol, and are involved in the endocytosis of various receptors and ligands with a multitude of mechanisms and regulation factors [31]. Ctx-B is a molecule that binds to glycosphingolipid GM1 on rafts to be subsequently internalized, so we used it as a marker for raft-mediated endocytosis [32]. Chlorpromazine and genistein were used in this study to inhibit clathrin- and raft-mediated endocytosis pathways, respectively.

Treatment with genistein was able to significantly decrease AgB oligomers uptake by RH and A549 cells. Accordingly, we found that AgB colocalizes with Ctx-B in RH cells. We observed a partial colocalization of AgB with Tfn and the inhibition assay with chlorpromazine showed only a slight, not significant, decrease in AgB internalization. It is possible that clathrin-mediated endocytosis accounts for just a small part of AgB uptake, making difficult the detection of a difference after inhibition by chlorpromazine. Taken together, our results are consistent with the idea that the endocytosis process is required for AgB entry into mammalian cells. Indeed, raft-mediated endocytosis is most likely the main pathway involved in AgB uptake by cells. However, a minor role for clathrin-mediated endocytosis in AgB internalization cannot be excluded.

It was proposed that AgB binds to macrophages and monocytes plasma membranes through a lipoprotein receptor; however no specific receptor could be determined [21]. Our findings are in agreement with this idea because some lipoprotein receptors, such as lectin-like oxidized LDL receptor-1 (LOX-1), LDL receptor-related protein 6 (LRP6), and scavenger receptors CD36 and CD204 use raft-mediated pathways for endocytosis [33–36]. Considering our results, it is also possible that more than one receptor might be involved in AgB binding, so that a higher efficiency of internalization is obtained. Alternatively, the receptor undergoes endocytosis by both pathways upon AgB binding. This regulatory mechanism involving different endocytic routes has been observed with LRP6, in which the receptor is internalized by caveolae (a raft subdomain) to promote Wnt/β-catenin signaling transduction, whereas the clathrin route leads to LRP6 degradation [37]. AgB uptake did not induce toxicity to cells according to our MTT assay, therefore internalization is more likely part of the mechanisms underlying AgB roles during *E. granulosus* metacestode infection. AgB-cell interaction may be a mechanism used by the parasite to create a more permissive microenvironment for metacestode development and survival. AgB presence in cytoplasm could interfere with cell metabolism, generating molecules and/or signals beneficial to the parasite.

The lack of genes coding for several key enzymes involved in fatty acid and cholesterol synthesis [15,16] reinforce the idea that AgB interaction with cells from the host tissue surrounding the hydatid cyst is a suitable scenario to get lipids from biological membranes or inner cell storages. A similar scenario has been described for the *Taenia solium* metacestode, where HLBPs were able to translocate lipid analogs to parasites’ tissues, and also colocalize with lipid droplets in the granuloma surrounding the metacestode [6]. Since lipids are essential for metacestode survival and development, the understanding of molecular mechanisms employed by the parasites to acquire these host macromolecules will provide potential targets for therapeutic discovery efforts.

AgB internalization by immune cells could influence the signalization towards anti-inflammatory or alternative pathways, eliciting a host non-protective response characteristic of immunoevasion processes. Similarly, in *Fasciola hepatica* a cathelicidin-like protein was described to bind lipid rafts, and after internalization, to divert macrophages function by suppressing lysosomal activity and, consequently, interfering with antigen presentation [38].

The destination of AgB after entry into cells was not evaluated here, but is an important issue to be addressed in further investigations in order to confirm that AgB internalization is necessary for its proposed biological roles. Among many possible destinations, an endocytosed protein can be routed to the late endosomes and lysosomes for degradation, to the trans-Golgi network or to recycling endosomes that bring the cargo back to the plasma membrane [39]. It would be interesting to know if AgB goes to the recycle pathway, which would permit its further exocytosis and return to parasitic tissues, or if it goes to lysosome for degradation. Further studies of colocalization with lysosomes and recycling endosomes using organelle markers described in the literature, as LAMP-1 and Rab-11 [40,41], should help to elucidate the fate of AgB after internalization.

Like AgB, cestode HLBPs are involved in parasite lipid homeostasis and immunological process [6,42]. Thus, further investigations on the cellular and molecular effect of HLBPs on host cells are important steps to improve the understanding of the parasites biology and disease progression. Likewise, elucidating how the molecules sequestered by HLBPs become available to parasites cells will help to identify potential targets for the treatment and control of cestodiasis.

## Acknowledgments

We thank the *Centro de Microscopia e Microanálise* (CMM-UFRGS) for technical support with confocal microscopy.

